# Diverse stresses converge on bacterial periplasmic expansion through distinct physiological mechanisms

**DOI:** 10.64898/2026.10.15.758610

**Authors:** Adrienne M. Brauer, Kevin Cho, Talia L. Shaw, Gary J. Patti, Kerwyn Casey Huang, Petra Anne Levin

## Abstract

Diverse environmental and antibiotic stresses induce localized periplasmic expansion–– the separation of the inner membrane from the cell wall and outer membrane––in Gram-negative bacteria, yet whether these morphologically similar responses arise through a common mechanism remains unclear. Here, we compare periplasmic expansion induced by sudden starvation, ciprofloxacin, mitomycin C, and globomycin in *Escherichia coli*. We find that these perturbations produce similar morphologies through distinct physiological and mechanical routes. Ciprofloxacin- and mitomycin C-induced periplasmic expansion coincides with loss of turgor, whereas starvation requires water efflux, and globomycin leads to unequal expansion of the inner and outer membrane as a result of aberrant inner membrane–cell wall crosslinking. Outer membrane stiffness modulates expansion during ciprofloxacin treatment. Despite these distinct mechanisms, robust periplasmic expansion during starvation and treatment with ciprofloxacin or globomycin requires UDP-glucose synthesis. Together, these findings establish periplasmic expansion as a convergent physical outcome of diverse stresses and reveal the remarkable plasticity of the Gram-negative cell envelope.

**Significance statement:** Localized periplasmic expansion is frequently visible in Gram-negative bacteria exposed to diverse stresses, yet it is rarely considered as a biological phenomenon in its own right. Consequently, the mechanisms underlying its formation and its relationship to bacterial physiology remain poorly understood. Here, we show that superficially similar periplasmic expansion can emerge through distinct physiological and mechanical routes during nutrient deprivation and antibiotic stress. Thus, rather than presenting as a single stereotyped stress response, periplasmic expansion is a convergent physical state that can arise from diverse perturbations. Our findings bring attention to a widespread but overlooked bacterial morphology that highlights the periplasm as a dynamic compartment and the Gram-negative cell envelope as a remarkably plastic cellular structure.

## Introduction

Bacteria undergo diverse morphological changes in response to environmental stress. Some, such as sporulation in *Bacillus subtilis,* fruiting body formation in *Myxococcus xanthus,* and heterocyst differentiation in *Anabaena*, are highly regulated developmental programs. Other morphological responses can emerge more directly from changes in cellular physiology and the physical state of the cell envelope. One such response is localized periplasmic expansion, in which the cytoplasm and inner membrane withdraw from the cell wall and outer membrane, resulting in an enlarged periplasmic region at a discrete subcellular location. In some cases, periplasmic expansion is accompanied by cytoplasmic condensation and increased cytoplasmic density^1,2^. Although striking when visualized, the physiological mechanisms underlying this morphology and its significance remain poorly understood.

The best-characterized example of periplasmic expansion is plasmolysis following hyperosmotic shock^3^, in which rapid water loss reduces turgor and causes differential contraction of the envelope layers^4^. However, periplasmic expansion also occurs independently of hyperosmotic shock. Related morphologies have been reported during nutrient limitation^1,5–8^, antibiotic exposure^1,2,9,10^, perturbation of envelope biogenesis^10–13^, and mutations affecting cellular signaling^14,15^ (**Table 1**). Periplasmic expansion can also occur during normal growth: in the deeply branching Gram-negative bacterium *Thermotoga maritima*, the outer membrane separates from the inner membrane at the cell poles in late exponential phase, producing its characteristic “toga.”^16^ Separation of envelope layers is not restricted to organisms with outer membranes, as related morphologies have also been observed in the Gram-positive bacterium *Bacillus subtilis* during kanamycin treatment^2^.

**Table 1:**
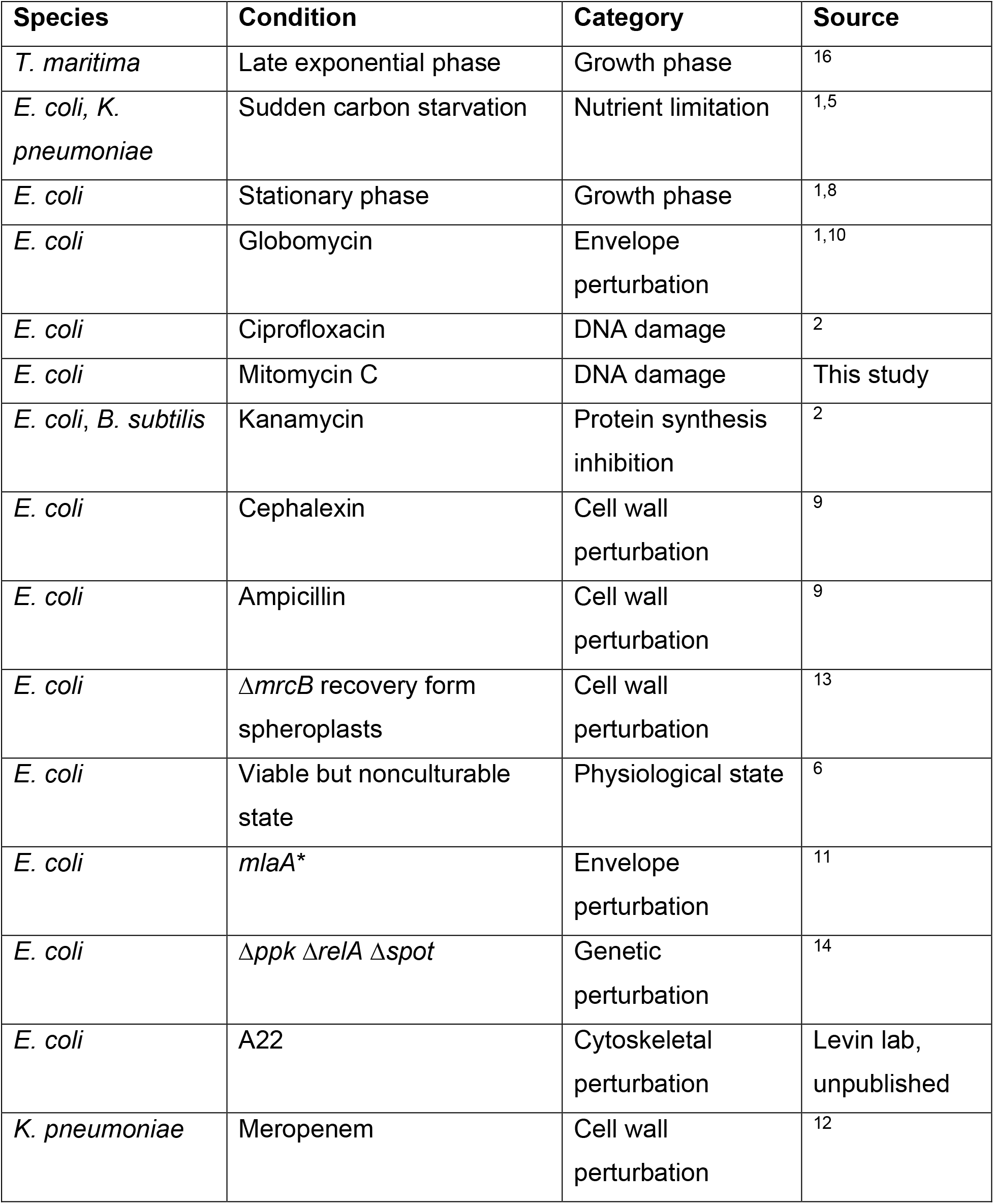
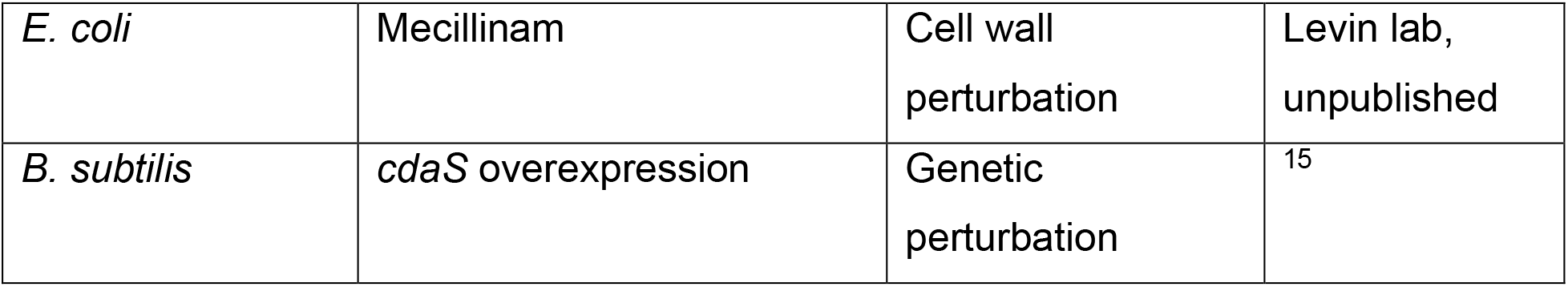
Conditions associated with periplasmic expansion.

The recurrence of periplasmic expansion under physiologically unrelated conditions raises a fundamental question: does this phenomenon represent a common cellular response to stress, or can distinct physiological perturbations converge independently on the same physical state? Our previous work identified one route to periplasmic expansion during sudden nutrient deprivation. Transfer of exponentially growing *Escherichia coli* to medium lacking carbon, nitrogen, or phosphate causes rapid cytoplasmic condensation and localized periplasmic expansion, predominantly at one pole^1^. Starvation-induced expansion is reversible and accompanied by loss of cytoplasmic water and increased cytoplasmic density, but occurs without a change in external osmolarity and is therefore distinct from classical plasmolysis^1,5^. It does not require new transcription or translation, canonical starvation-response pathways, or major mechanosensitive channels, suggesting that it instead reflects a rapid physical response involving water redistribution and mechanical coupling among the layers of the Gram-negative cell envelope.

A morphologically similar response occurs following antibiotic treatment, despite a very different initiating perturbation. Ciprofloxacin, a fluoroquinolone that inhibits DNA gyrase, also induces cytoplasmic condensation and periplasmic expansion. Reduced turgor pressure in these cells suggests a turgor loss-dependent mechanism ^2^. Thus, starvation and ciprofloxacin produce superficially similar envelope morphologies despite differing origins. Whether these and other forms of periplasmic expansion arise through a shared physiological pathway or instead represent independent routes to a common physical state remains unknown.

To determine whether periplasmic expansion represents a common physiological response or a convergent physical outcome, we systematically compared its dynamics, physiology, and mechanical determinants across four distinct perturbations: sudden starvation, ciprofloxacin, mitomycin C, and globomycin. We identify both stress-specific determinants and shared requirements, demonstrating that visually similar periplasmic expansion can emerge through distinct physiological and mechanical routes. Together, our findings establish periplasmic expansion as a convergent physical state of the Gram-negative cell envelope and reveal the periplasm as a dynamic compartment whose dimensions can respond dramatically to changes in cellular physiology.

## Results

### Periplasmic expansion exhibits stress-dependent morphology, dynamics, and physiological outcomes

To determine whether periplasmic expansion induced by different stresses represents a physiologically equivalent cellular state, we examined exponentially growing *E. coli* MG1655 following sudden starvation^1^ or treatment with three antibiotics with distinct cellular targets: ciprofloxacin^2^, which inhibits DNA gyrase; mitomycin C, which damages DNA; and globomycin^1^, which inhibits maturation of Lpp, the abundant lipoprotein that links the outer membrane to the cell wall^17^ (**Methods**).

We first asked whether the phase-bright regions observed under each condition reflected separation of the inner membrane from the outer envelope, as previously established during sudden starvation^1^. Under all four conditions, cytoplasmic GFP was excluded from phase-bright regions that accumulated periplasmic mCherry (**Fig. 1A**), while HADA-labeled cell wall remained exterior to the cytoplasm (**Fig. S1A**). No expanded regions containing periplasmic mCherry were observed in unstressed exponentially growing cells (**Fig. S1A,B**). Thus, despite their distinct cellular targets, all four perturbations induced separation of the cytoplasm and inner membrane from the outer layers of the cell envelope.

**Figure 1:**
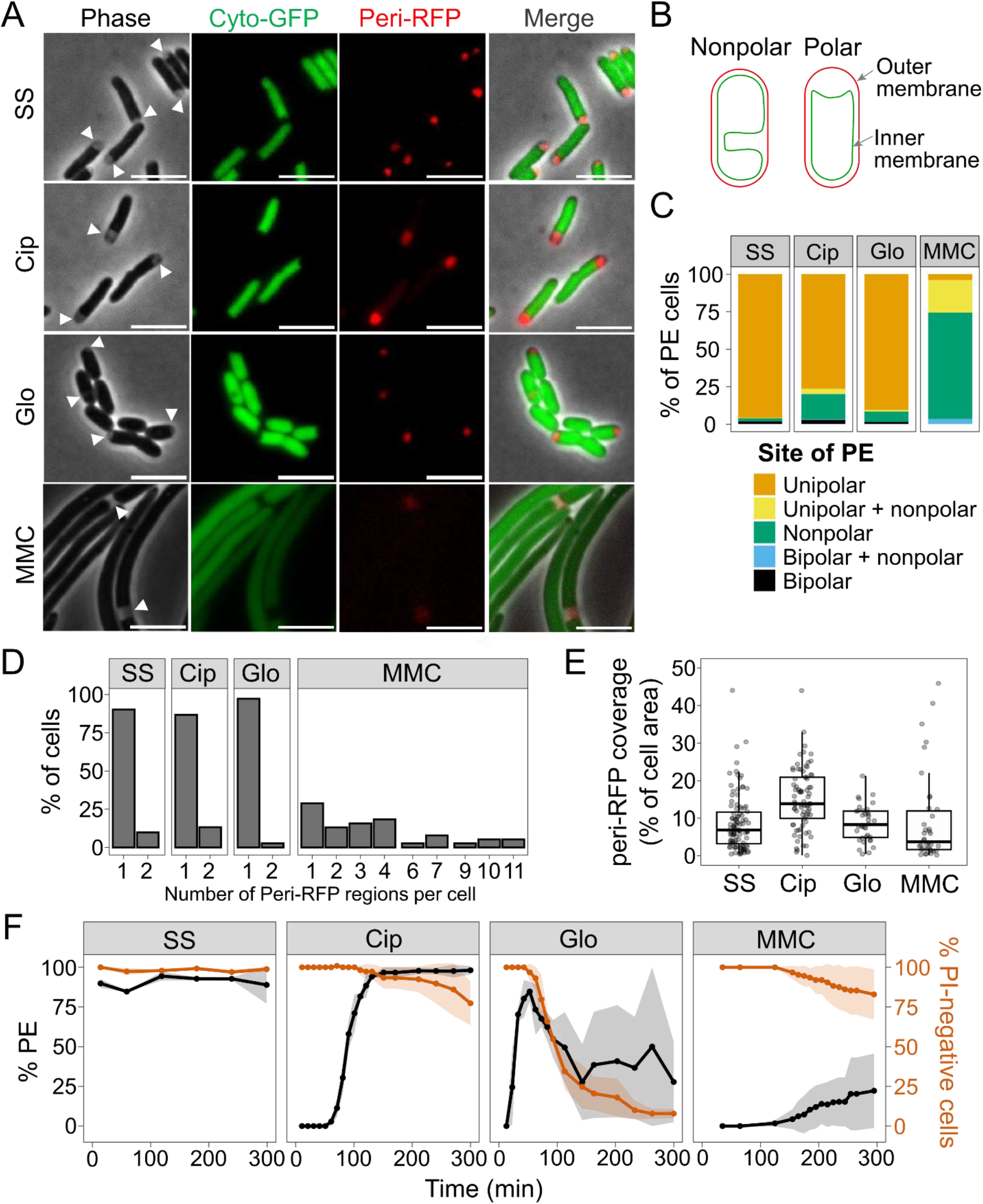
Distinct stresses induce periplasmic expansion with different morphologies, dynamics, and physiological outcomes. Cells expressing an IPTG-inducible mCherry (RFP) protein fused to a periplasmic signal sequence and a constitutive GFP were grown to log phase then exposed to sudden starvation (SS), ciprofloxacin (Cip), globomycin (Glo), or mitomycin C (MMC). Chloramphenicol and IPTG were maintained in the media throughout for selection and induction, respectively. **A**, Representative phase-contrast and fluorescence images of cells expressing cytoplasmic GFP (Cyto-GFP) and periplasmic mCherry (Peri-RFP). Arrowheads indicate regions of periplasmic expansion (PE). Scale bars, 5 µm. **B**, Schematic illustrating polar and nonpolar PE. **C**, Mean percentage of cells exhibiting PE at unipolar, bipolar, nonpolar, or combined polar and nonpolar sites. **D**, Distribution of the number of discrete Peri-RFP regions per cell exhibiting PE. **E**, Fraction of total cell area occupied by Peri-RFP regions in cells exhibiting PE. For cells containing multiple regions, their areas were summed. Boxes indicate the interquartile range (IQR) and median; whiskers extend to 1.5× IQR. **F**, Percentage of cells exhibiting PE among PI-negative cells (black, left axis) and percentage of cells remaining PI-negative relative to the initial cell population (orange, right axis) following initiation of each stress. Lines indicate means and shaded regions indicate SD.

The extent and morphology of periplasmic expansion nevertheless differed substantially among stresses. Expansion predominantly occurred as a single discrete region at one pole during starvation and treatment with ciprofloxacin or globomycin (**Fig. 1B-D, Supplementary Table 1**). In contrast, mitomycin C treatment caused extensive filamentation (**Fig. S1C**), consistent with division inhibition during the SOS response^18^, and produced numerous small nonpolar periplasmic expansions along the cell envelope. These regions were most readily detected by accumulation of periplasmic mCherry, with only slight indentations apparent by phase-contrast microscopy (**Fig. S1D**). The expanded periplasm occupied ∼7% of cell volume following sudden starvation, ∼16% following ciprofloxacin treatment, ∼8% following globomycin treatment, and ∼4% following mitomycin C treatment (**Fig. 1D**). Fluorescent protein overexpression also appeared to generate inactive aggregates^19^ following mitomycin C treatment that were visible as phase-bright regions and excluded cytoplasmic GFP but did not accumulate periplasmic mCherry (**Fig. S1E**). We therefore used phase-contrast microscopy to score periplasmic expansion in subsequent experiments, avoiding potential artifacts associated with fluorescent protein expression.

The kinetics of periplasmic expansion also varied substantially across conditions. (**Fig. 1F**). Expansion occurred rapidly, within 10 min of sudden starvation^1^, and within ∼20 min of globomycin treatment, whereas it emerged only after ∼1 h of ciprofloxacin treatment. Mitomycin C produced the slowest response, with fewer than 25% of cells exhibiting periplasmic expansion after 5 h. Because starved cells underwent reductive division and reduced the degree of periplasmic expansion within ∼20 min on agarose pads, we quantified starvation time courses by sampling cells directly from liquid culture.

We next asked whether periplasmic expansion was associated with loss of membrane integrity, as measured by uptake of propidium iodide (PI). Suddenly starved cells^1^ and cells treated with ciprofloxacin or mitomycin C remained predominantly PI-negative for >5 h (99±2%, 77±14%, and 83±16% respectively; **Fig. 1F**). In contrast, 92±3% of globomycin-treated cells took up PI within 5 h. Among ciprofloxacin- and mitomycin C-treated cells that ultimately became PI-positive, PI uptake typically occurred 1–3 h after the onset of periplasmic expansion (**Fig. S1F**). Single-cell dynamics could not be resolved during sudden starvation because of the sampling method, but the low overall frequency of PI uptake similarly argues against a close association between expansion and membrane permeabilization. Together, these results establish periplasmic expansion as a common morphological outcome of diverse stresses whose geometry, kinetics, and relationship to membrane integrity vary substantially with the inducing perturbation.

### Periplasmic expansion can occur with or without detectable turgor loss

Previous work showed that ciprofloxacin-treated cells exhibiting periplasmic expansion have lost turgor pressure^2^. Loss of turgor causes a decrease in cell length, the magnitude of which is influenced by outer membrane stiffness^4^, and can produce periplasmic expansion during plasmolysis. We therefore tracked single-cell length dynamics before and during periplasmic expansion to determine whether turgor loss accompanies expansion across stress conditions and whether the two events are temporally associated.

Ciprofloxacin- and mitomycin C-treated cells abruptly decreased in length by 9.5±1.8% and 8.4±4.0%, respectively, coincident with the onset of periplasmic expansion (**Fig. 2A-C**). These decreases were comparable in magnitude to those produced by osmotic perturbations that eliminate turgor^4^ (**Fig. S2A**), indicating that periplasmic expansion under both conditions is associated with loss of turgor. In a small subset of mitomycin C-treated cells (<1%), the inner membrane subsequently re-expanded and growth resumed, indicating that the physiological state associated with expansion can be reversible (**Fig. S2B**). By contrast, suddenly starved cells exhibited no detectable decrease in length at the onset of periplasmic expansion, while globomycin-treated cells continued to elongate (**Fig. 2B**). Thus, detectable whole-cell turgor loss accompanies periplasmic expansion under some stress conditions but is not required for the phenotype.

**Figure 2:**
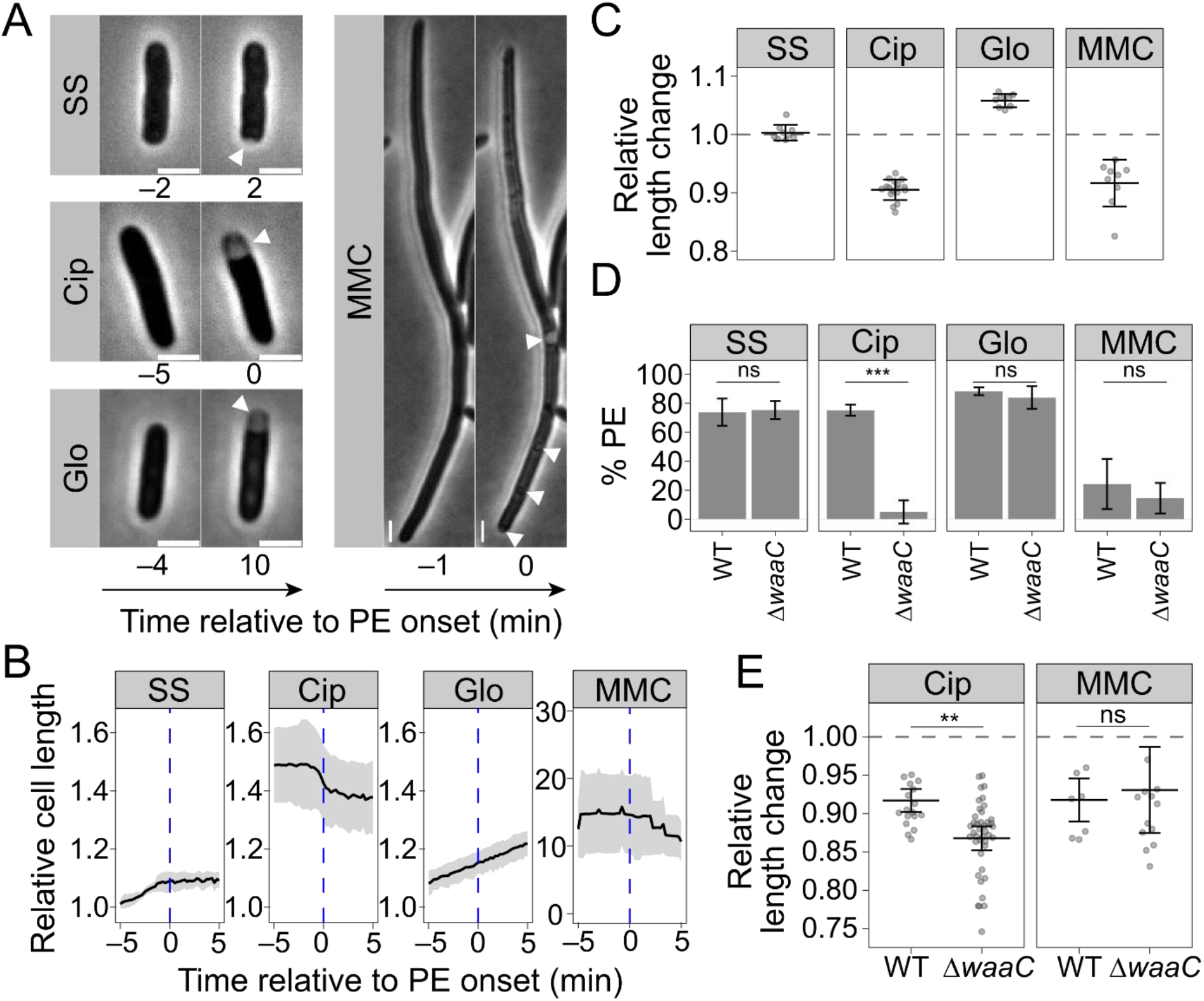
Periplasmic expansion can occur with or without detectable turgor loss. **A**, Representative phase-contrast images of cells immediately before and after the onset of periplasmic expansion (PE) during sudden starvation (SS) or treatment with ciprofloxacin (Cip), globomycin (Glo), or mitomycin C (MMC). Numbers indicate time relative to PE onset. Arrowheads indicate sites of PE. Scale bars, 2 µm. **B**, Relative cell length over time, aligned to the onset of PE (blue dashed line). Lines indicate means and shaded regions indicate SD. **C**, Relative cell length following PE compared with cell length before PE for the cells analyzed in **B**. Points represent individual cells; bars indicate mean ± SD. **D**, Percentage of PI-negative BW25113 (WT) and Δ*waaC* cells exhibiting PE following the indicated treatments. Bars indicate mean ± SD. Statistical significance was determined using a two-tailed Student’s t-test. ns, not significant (*p* ≥ 0.05); ***: *p* < 0.001. **E,** Relative cell length following PE in ciprofloxacin- and MMC-treated WT cells and following the corresponding abrupt decrease in cell length in Δ*waaC* cells. Points represent individual cells from three biological replicate experiments; bars indicate the mean of replicate means ± SD. Statistical significance was determined using a linear mixed-effects model (LMM) accounting for biological replicate as a random effect, followed by Dunnett’s post-hoc test for pairwise comparisons against the WT control within each condition. ns, not significant (*p* ≥ 0.05); **: *p* < 0.01.

We next tested whether the mechanical properties of the outer envelope determine whether turgor loss is manifested as visible periplasmic expansion. Specifically, we reasoned that if the outer membrane and cell wall resist contraction as the cytoplasm and inner membrane shrink, reducing outer membrane stiffness should allow greater contraction of the outer envelope and thereby reduce the separation between envelope layers. To test this model, we examined Δ*waaC* cells, which have a truncated LPS core and substantially reduced outer membrane stiffness^20^. Consistent with a requirement for a stiff outer membrane, ciprofloxacin-treated Δ*waaC* cells exhibited almost no visible periplasmic expansion, in contrast to the BW25113 parent (**Fig. 2D**). Δ*waaC* cells decreased in length by 13.2±1.6%, 59% more than the parent strain (**Fig. 2E**), indicating that turgor loss still occurred. This is consistent with our prediction that a higher degree of outer membrane contraction could reduce the visible separation between envelope layers.

Unexpectedly, Δ*waaC* did not alter either the fraction of cells exhibiting periplasmic expansion or the magnitude of cell shortening following mitomycin C treatment (**Fig. 2D,E**), indicating that turgor loss alone is insufficient to determine how the envelope deforms. Mitomycin C-treated Δ*waaC* cells also exhibited frequent oscillations in cell length that were absent from the parent strain (**Fig. S2C**), suggesting that additional properties of the envelope influence its mechanical response under this condition. Starvation- and globomycin-induced expansion likewise occurred at near wild-type levels in Δ*waaC* cells (**Fig. 2D**), consistent with their occurrence in the absence of detectable whole-cell turgor loss.

Together, these results distinguish at least two physiological states associated with periplasmic expansion. Ciprofloxacin- and mitomycin C-induced expansion coincides with detectable turgor loss, whereas starvation- and globomycin-induced expansion does not. Moreover, the Δ*waaC* phenotype demonstrates that turgor loss alone is insufficient to produce visible separation of the envelope layers and that outer membrane mechanics can determine how changes in cellular pressure are translated into envelope morphology.

### Globomycin-mediated aberrant Lpp crosslinking leads to unequal expansion of the inner and outer membranes

Globomycin is unique among the four conditions tested in that it directly perturbs mechanical coupling between layers of the envelope through its effects on Lpp^17^. Lpp is the most abundant protein in *E. coli*^21^ and provides the major covalent linkage between the outer membrane and peptidoglycan^22,23^, thereby contributing substantially to envelope mechanics^4,20,24^. Globomycin inhibits Lpp trafficking^25^, trapping newly synthesized Lpp in the inner membrane and enabling its aberrant crosslinking to peptidoglycan^17,26^.

Globomycin-treated cells continued to grow while exhibiting periplasmic expansion, unlike cells subjected to sudden starvation or treatment with ciprofloxacin or mitomycin C (**Fig. 2B**). We therefore hypothesized that aberrant crosslinking between the inner membrane and peptidoglycan disrupts coordinated expansion of the envelope during growth, resulting in localized separation of its layers. Consistent with this model, periplasmic expansion was essentially abolished in both *Δlpp* and Δ*ldtABCDEF* cells, decreasing by 100% and 95%, respectively (**Fig. 3A, S3A**). Neither mutation affected periplasmic expansion during sudden starvation or mitomycin C treatment (**Fig. 3A**). Both mutations also reduced the frequency of ciprofloxacin-induced expansion by 50–60%, although the magnitude of ciprofloxacin-induced cell shortening was unaffected (**Fig. S3B**).

**Figure 3:**
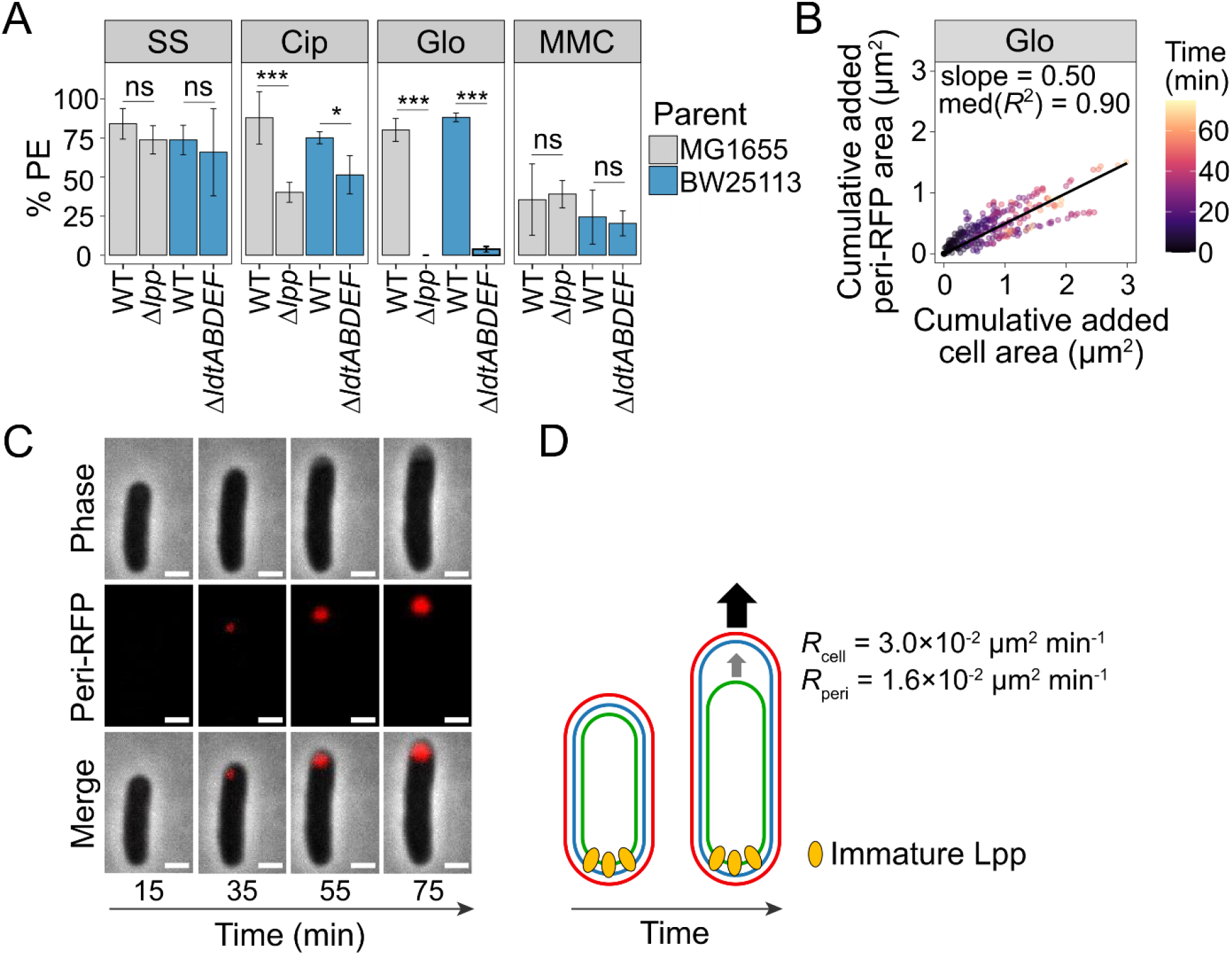
Globomycin-induced periplasmic expansion requires aberrant Lpp crosslinking and cell growth. **A**, Percentage of Δ*lpp*, Δ*ldtABCDEF*, and their corresponding wild-type (WT) parent cells, MG1655 or BW25113, exhibiting periplasmic expansion (PE) following sudden starvation (SS) or treatment with ciprofloxacin (Cip), globomycin (Glo), or mitomycin C (MMC). BW25113 WT data is replotted from Fig. 2A for statistical comparisons. Bars indicate mean ± SD. Statistical significance was determined using a one-way ANOVA followed by Dunnett’s multiple-comparison test. ns, not significant (*p* ≥ 0.05); **, *p* < 0.01; ***, *p* < 0.001. **B**, Cumulative increase in periplasmic area plotted against cumulative increase in total cell area for individual globomycin-treated cells, relative to the first time point at which a discrete periplasmic mCherry region was detected (*t* = 0). Points are colored according to time after PE onset. Individual linear regressions were fitted to cells tracked for ≥ 4 frames after PE onset. The black line indicates the population mean expansion trajectory, calculated from the mean slope of individual linear regressions fit to each cell, constrained to pass through the origin (0,0); slope = 0.50 and median *R*^2^ = 0.90. **C**, Representative time-lapse images of a globomycin-treated cell showing progressive increases in cell area and the area of the expanded periplasm. Numbers indicate time after globomycin addition. Scale bars, 2 µm. **D**, Model in which aberrant crosslinking of immature Lpp between the inner membrane and peptidoglycan locally restricts coordinated expansion of the envelope during growth, resulting in PE at a distal site. Representative rates of increase in total cell area (*R*_cell_) and periplasmic area (*R*_peri_) are indicated.

To determine how periplasmic expansion develops as globomycin-treated cells grow, we tracked periplasmic mCherry fluorescence over time. The area of the expanded periplasm increased approximately linearly (median *R*^2^ = 0.90; **Fig. 3B,C**) and at approximately half the rate of increase in whole-cell area (slope = 0.496). Thus, periplasmic expansion progressively increased as cells grew, consistent with differential expansion of the envelope layers. Supporting a requirement for ongoing growth, treatment with either rifampicin or chloramphenicol, which inhibits RNA or protein synthesis, respectively, reduced the frequency of globomycin-induced periplasmic expansion by 95% (**Fig. S3C,D**).

Because Lpp cannot crosslink the inner membrane to peptidoglycan within the expanded region itself, the aberrant crosslinks responsible for expansion must act elsewhere in the cell. We therefore propose a model in which aberrant Lpp crosslinking locally anchors the inner membrane to peptidoglycan, preventing coordinated expansion of the inner and outer envelope as the cell grows and producing periplasmic expansion at a distal site (**Fig. 3D**). Together, these results support a growth-dependent mechanism for globomycin-induced periplasmic expansion in which aberrant Lpp crosslinking disrupts coordinated expansion of the envelope layers.

### Hypoosmotic shift reverses starvation-induced periplasmic expansion

Unlike turgor loss, which is accompanied by contraction of the load-bearing outer envelope, sudden starvation-induced loss of cytoplasmic water occurs without detectable whole-cell contraction (**Fig. 2B**), suggesting redistribution of water between envelope compartments rather than loss of whole-cell turgor. We therefore asked whether reducing external osmolarity could reverse starvation-induced periplasmic expansion. Following sudden starvation, cells exhibiting robust periplasmic expansion were subjected to an additional wash in 0.5X M9 salts. Reducing external osmolarity eliminated 99% of the observed periplasmic expansion (**Fig. 4A,B**). By contrast, shifting cells to 0.5X LB had little effect on periplasmic expansion induced by ciprofloxacin, mitomycin C, or globomycin (**Fig. 4B**). Thus, starvation-induced periplasmic expansion is particularly sensitive to external osmolarity, consistent with redistribution of water across the cell envelope underlying this response.

**Figure 4:**
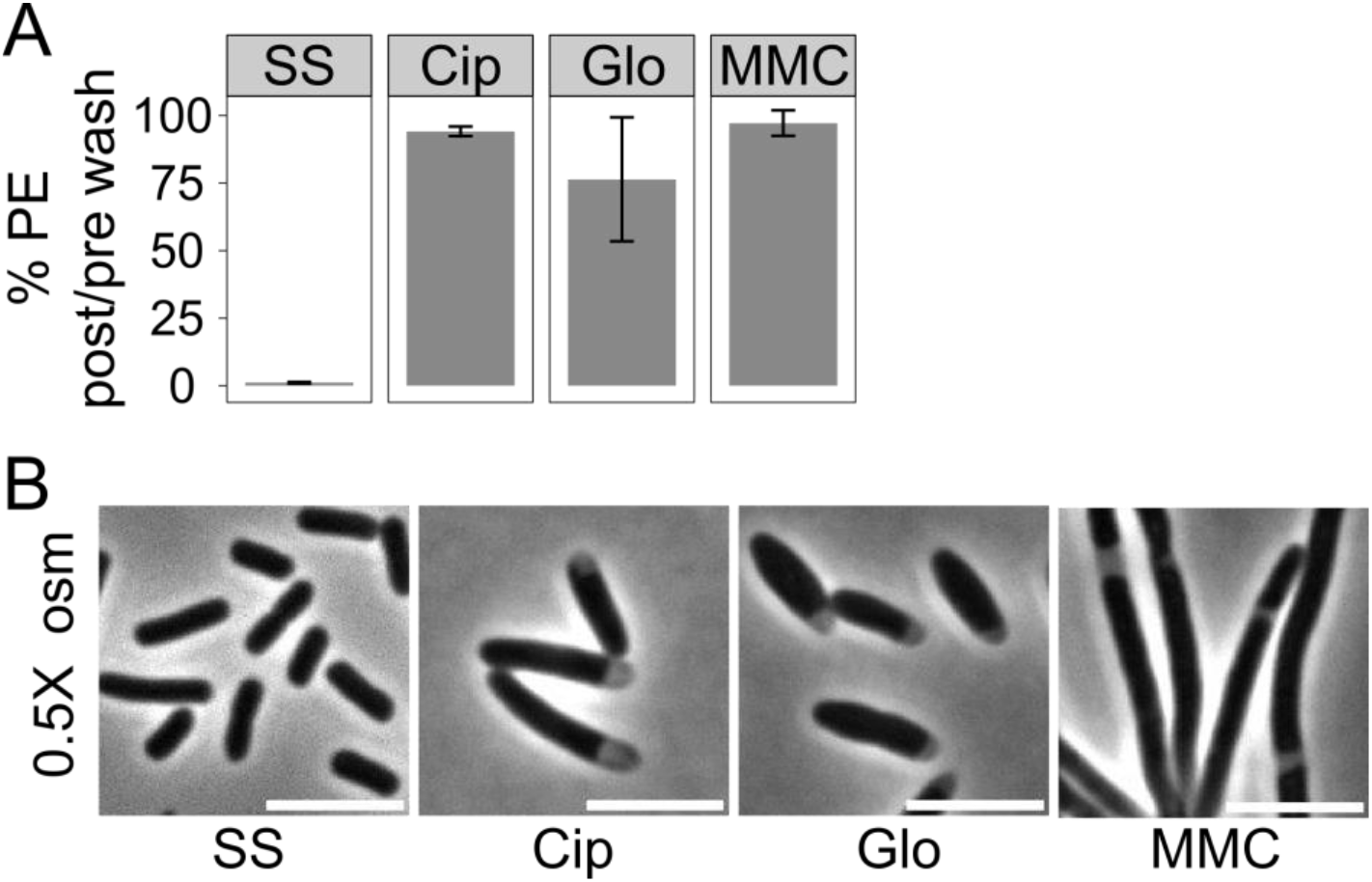
Hypoosmotic shift reverses starvation-induced periplasmic expansion. **A**, Cells were subjected to sudden starvation (SS) or treated with ciprofloxacin (Cip), globomycin (Glo), or mitomycin C (MMC) for 5, 100, 50, or 180 min, respectively, before exposure to a hypoosmotic shift. Starved cells were washed with 0.5X M9 salts, whereas antibiotic-treated cells were washed with 0.5X LB containing the respective antibiotic. The percentage of cells exhibiting periplasmic expansion (PE) after the hyperosmotic shift is shown relative to the percentage exhibiting PE immediately before the shift. Bars indicate mean ± SD. **B**, Representative phase-contrast images following the hypoosmotic shift. Scale bars, 5 µm.

### UDP-glucose synthesis is required for robust periplasmic expansion across distinct stresses

Given the contributions of osmotic and mechanical properties of the envelope to periplasmic expansion, we asked whether UDP-glucose synthesis, which contributes to multiple envelope-associated pathways, influences expansion across stress conditions. UDP-glucose is a precursor for osmoregulated periplasmic glucans (OPGs), which modulate envelope physiology in response to osmotic conditions^27^, as well as the LPS core oligosaccharide and other cellular products (**Fig. S4A**).

UDP-glucose is synthesized from glucose-6-phosphate in two steps (**Fig. 5A**). Pgm interconverts glucose-6-phosphate and glucose-1-phosphate, and GalU converts glucose-1-phosphate and UTP to UDP-glucose. Deletion of *galU* reduced the frequency of periplasmic expansion by 99% following sudden starvation and by more than 80% following ciprofloxacin and globomycin treatment, whereas mitomycin C-induced expansion was unaffected (**Fig. 5B**). Deletion of *pgm* similarly reduced expansion under these three conditions, further supporting a requirement for UDP-glucose synthesis^28^. Catalytically inactive GalU failed to complement complement the Δ*galU* phenotype (**Fig. S4B**), demonstrating that GalU enzymatic activity is required. However, mutations disrupting single or multiple downstream UDP-glucose-dependent pathways did not recapitulate the expansion defect of Δ*galU* (**Fig. S4C**), suggesting that the phenotype cannot be attributed to loss of any downstream pathway tested.

**Figure 5:**
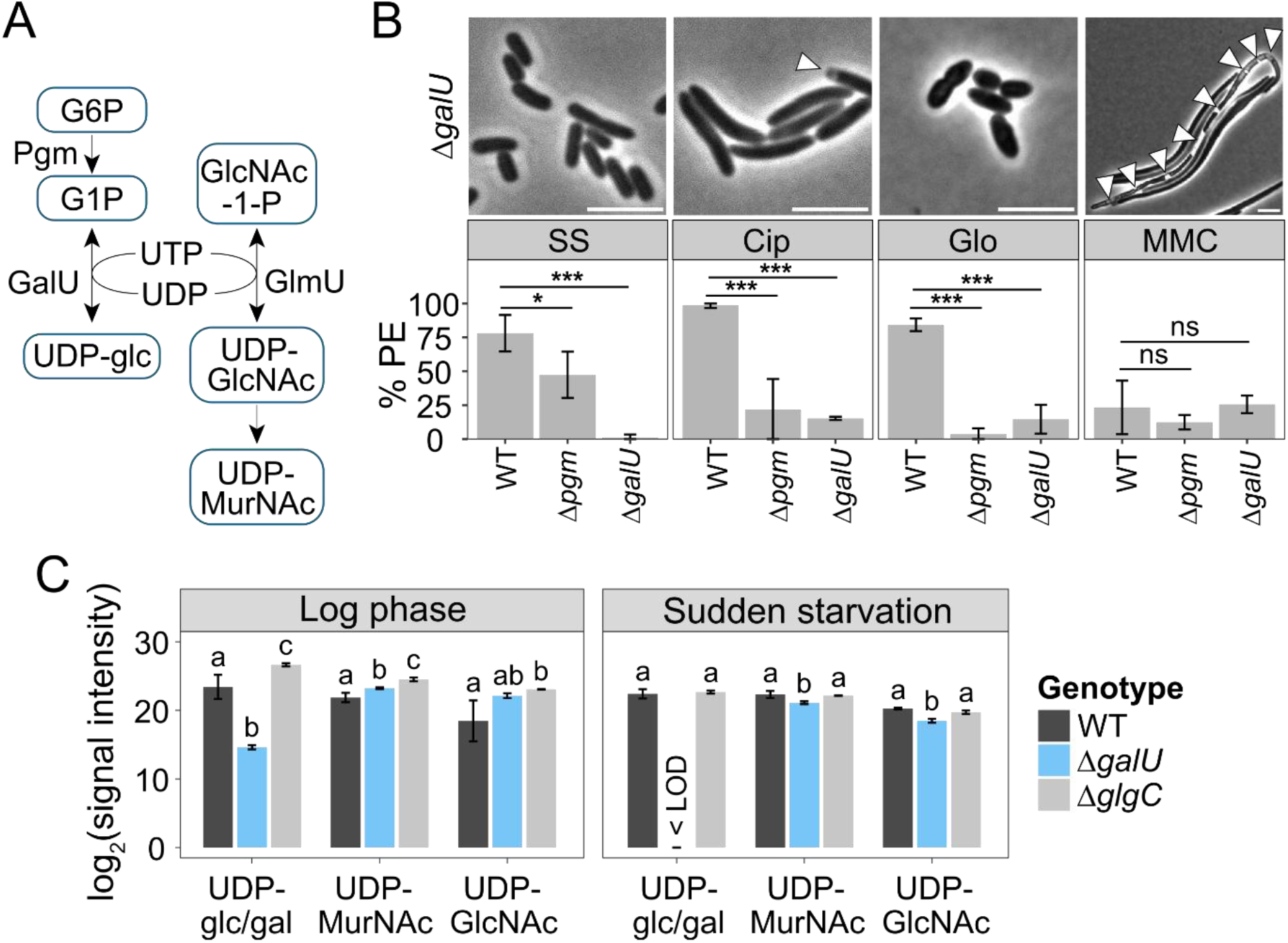
UDP-glucose synthesis is required for robust periplasmic expansion across distinct stresses. **A**, Schematic of UDP-glucose (UDP-glc), UDP-*N*-acetyl-glucosamine (UDP-GlcNAc), and UDP-*N*-acetyl-muramic acid (UDP-MurNAc) synthesis. **B**, Representative phase-contrast images of Δ*galU* cells following sudden starvation (SS) or after treatment with ciprofloxacin (Cip), globomycin (Glo), or mitomycin C (MMC) (top). Scale bars, 5 µm. Arrowheads indicate regions of periplasmic expansion (PE). Percentage of wild-type (WT), Δ*galU*, and Δ*pgm* cells exhibiting PE under each condition (bottom). MG1655 WT data is replotted from Fig. 3A for statistical comparisons. Bars indicate mean ± SD. Statistical significance relative to WT within each condition was determined using a one-way ANOVA followed by Dunnett’s multiple-comparison test. ns, not significant (*p* ≥ 0.05); *, *p* < 0.05; ***, *p* < 0.001. **C**, Relative abundance (log_2_ transformed signal intensity) of the indicated nucleotide sugars measured by LC-MS in WT, Δ*galU*, and Δ*glgC* cells during exponential growth and 45 min after sudden starvation. Bars indicate mean ± SD. <LOD, below the limit of detection. For each metabolite and growth condition, differences among genotypes were evaluated by one-way ANOVA followed by Tukey’s HSD test (α = 0.05). Letters (a, b, c) represent statistical significance within each metabolite and condition; genotypes sharing a letter are not significantly different (*p* ≥ 0.05).

We therefore asked whether loss of *galU* produces broader metabolic changes that might distinguish Δ*galU* cells from mutants that retain periplasmic expansion. We performed metabolomics on wild-type, Δ*galU*, and Δ*glgC* cells during exponential growth and 45 min after sudden starvation. Δ*glgC* cells, which are defective in the first committed step of glycogen synthesis but exhibit wild-type levels of periplasmic expansion (**Fig. S4A,C**), provided a comparison for perturbation of glucose-1-phosphate metabolism without loss of expansion. As expected, the combined signal from the indistinguishable isomers UDP-glucose and UDP-galactose was approximately 1,000-fold lower in exponentially growing *ΔgalU* cells than in wild type and fell below the limit of detection following starvation **(Fig. 4C**).

Despite this profound depletion, Δ*galU* cells did not exhibit a globally distinct metabolic state. During exponential growth, the Δ*galU* metabolomic profile clustered closely with that of Δ*glgC* cells by principal-component analysis, and all three genotypes converged after starvation (**Fig. S4D**). We therefore identified metabolites whose abundance in starved Δ*galU* cells differed most strongly from both wild-type and Δ*glgC* cells. Notably, UDP-*N*-acetylglucosamine (UDP-GlcNAc) and UDP-*N*-acetylmuramic acid (UDP-MurNAc), two nucleotide sugars required for envelope biosynthesis, ranked among the top 15 most strongly altered metabolites (**Fig. 4A, Fig. S4E,F**). Their abundance was intermediate between wild-type and Δ*glgC* cells during exponential growth but fell to less than half the levels observed in both strains following starvation (**Fig. 5C**). Thus, loss of GalU perturbs not only UDP-glucose abundance but also other nucleotide-sugar pools, particularly during starvation. Together, these results identify UDP-glucose synthesis as a shared requirement for robust periplasmic expansion induced through three otherwise distinct physiological routes, suggesting that its role may reflect broader coupling within nucleotide-sugar metabolism rather than production of a single UDP-glucose-derived product.

## Discussion

Our findings establish periplasmic expansion as a convergent morphological outcome that can emerge from distinct physiological and mechanical perturbations. During sudden starvation, cytoplasmic water loss and periplasmic expansion occur without detectable whole-cell contraction, whereas expansion during ciprofloxacin and mitomycin C treatment coincides with loss of turgor. The inability of a hypoosmotic shift to reverse expansion in ciprofloxacin- and mitomycin C-treated cells further distinguishes these states from starvation-induced expansion (**Fig. 4**). One possible explanation is the membrane-permeabilization model proposed previously for ciprofloxacin treatment^2^, in which loss of small osmolytes prevents restoration of turgor even when external osmolarity is reduced. The persistence of PI exclusion during the initial expansion indicates that any such permeability change precedes gross loss of membrane integrity. Globomycin induces expansion through a third route requiring aberrant covalent crosslinking between the inner membrane and peptidoglycan and continued cell growth (**Fig. 3**). Together, these findings demonstrate that similar separation of envelope layers can emerge from changes in cellular pressure, water distribution, or mechanical coupling during growth.

The physiological consequences of periplasmic expansion remain less clear. Starved cells maintain membrane integrity and viability despite extensive expansion, and the phenotype is rapidly reversible upon nutrient restoration^1^. Similarly, a subset of mitomycin C-treated cells re-expanded their inner membranes and resumed growth (**Fig. S2b**), demonstrating that substantial separation of the envelope layers is not necessarily a terminal state. Whether expansion itself promotes survival remains unknown. In conditions associated with turgor loss, the accompanying cessation of growth (**Fig. 2b**) could potentially increase tolerance to antibiotic stress^29^, whereas in other contexts disruption of normal spatial relationships between envelope layers could instead be detrimental.

A notable point of convergence among otherwise distinct mechanisms is their dependence on UDP-glucose synthesis. Loss of *galU* or *pgm* strongly reduced periplasmic expansion during starvation and treatment with ciprofloxacin and globomycin, but not mitomycin C treatment. The failure of downstream pathway mutants to phenocopy Δ*galU* argues against a requirement for the known UDP-glucose-derived products tested. Instead, metabolomic analysis revealed broader changes in nucleotide-sugar metabolism in Δ*galU* cells, including reduced UDP-GlcNAc and UDP-MurNAc following starvation. Because these metabolites contribute to envelope biosynthesis, these changes raise the possibility that UDP-glucose synthesis influences periplasmic expansion indirectly through the metabolic state of the envelope. Determining how nucleotide-sugar metabolism influences the physical relationships among envelope layers will require further investigation.

Periplasmic expansion is also likely to alter processes that depend on the normal spatial organization of the cell envelope. Increasing periplasmic volume should reduce the concentrations of soluble periplasmic components while increasing the distance between the inner membrane, cell wall, and outer membrane. Many processes–– including cell wall synthesis, outer-membrane biogenesis, nutrient transport, and retrograde envelope signaling––depend on protein complexes or molecular interactions that bridge these layers. Consistent with the importance of this geometry, experimentally increasing periplasmic width can impair signaling from the envelope to the cytoplasm^30^. Localized expansion could therefore spatially restrict envelope synthesis, transport, or signaling and, in some contexts, increase rather than decrease vulnerability to stress. Conversely, the spatially restricted nature of periplasmic expansion could provide a useful means of probing processes that depend on interactions between envelope layers, particularly at the cell poles, where putative anterograde phospholipid transporters have been reported to localize^31^. The rapid reversal of expansion following starvation^1^ and in a subset of mitomycin C-treated cells (**Fig. S2B**) further demonstrates that these spatial relationships can be substantially disrupted and subsequently restored.

More broadly, our findings highlight the periplasm as a mechanically and osmotically dynamic cellular compartment rather than a fixed space between the two membranes. The loss of cytoplasmic volume during starvation without detectable whole-cell contraction suggests that pressure can be redistributed across the envelope, potentially allowing increased periplasmic osmotic pressure to maintain tension on the peptidoglycan-outer membrane complex as cytoplasmic volume decreases. Such a model is consistent with recent evidence that the Gram-negative periplasm can sustain substantial osmotic pressure^24^. In this view, changes in periplasmic volume provide a means of redistributing water and mechanical stresses among the layers of the envelope without necessarily changing overall cellular dimensions. Whether periplasmic expansion is adaptive in particular contexts or simply an emergent physical consequence of stress remains unresolved. Regardless, its occurrence across diverse perturbations reveals an unexpected degree of plasticity in bacterial cellular architecture and establishes the periplasm as an important component of the physical response of cells to environmental change.

## Materials and Methods

### Bacterial strains, plasmids, and growth conditions

Strains and plasmids used in this study are listed in **Table S3**. Unless otherwise noted, deletion alleles were transferred from the Keio collection into the *E. coli* MG1655 background by P1 transduction. The Δ*waaC* mutant was examined in the BW25113 background because the *waaC* deletion prevented P1 transduction into MG1655. The Δ*ldtABCDEF* strain was used in its original background because of the absence of selectable markers for transfer by P1 transduction. Where indicated, kanamycin-resistance cassettes were removed by FLP-mediated recombination to generate markerless alleles. The *galU*(D137N) allele was generated using *In Vitro* Assembly (IVA)^32^.

Unless otherwise noted, cells were cultured in lysogeny broth (LB; 10 g/L tryptone, 10 g/L NaCl, and 5 g/L yeast extract) at 37 °C with shaking at 200 rpm. Where appropriate, antibiotics or inducers were added at the following concentrations: chloramphenicol, 30 µg/mL; IPTG, 100 µM. M9 salts were prepared by diluting a 5X stock (64 g/L Na_2_HPO_4_•7H_2_O, 15 g/L KH_2_PO_4_, 2.5 g/L NaCl, and 5 g/L NH_4_Cl) to 1X and supplementing with 1 mM MgSO_4_ and 0.1 mM CaCl_2_. M9 salts used for starvation contained no added carbon source.

### Assay for periplasmic expansion and propidium iodide uptake

Cells were grown to mid-exponential phase, diluted to an OD_600_ of 0.005, and harvested upon reaching early exponential phase (OD_600_ = 0.1–0.2). For antibiotic-induced periplasmic expansion, cells were spotted onto LB pads containing 1% agarose, 1.5 µM PI, and either 10 µg/mL ciprofloxacin, 20 µg/mL globomycin, or 18 µg/mL mitomycin C. When indicated, 30 µg/mL chloramphenicol or 50 µg/mL rifampicin was included to inhibit growth.

For starvation experiments, cells were pelleted at 6,010*g* for 1 min and washed three times with 1X M9 salts, as described previously^1^. Cells were then spotted onto 1% agarose pads prepared with 1X M9 salts and supplemented with 1.5 µM PI.

For hypoosmotic-shift experiments, antibiotic-treated cells were exposed to globomycin, ciprofloxacin, or mitomycin C in liquid culture for 50, 100, or 180 minutes, respectively, before being pelleted and washed with 0.5X LB containing the same concentration of the respective antibiotic. Starved cells were similarly pelleted and washed with 0.5X M9 salts. Cells were spotted onto 1% agarose pads made with 0.5X LB or 0.5X M9 salts and imaged immediately.

Cells exhibiting periplasmic expansion were identified by phase-contrast microscopy based on the presence of visible phase bright regions. The frequency of periplasmic expansion was calculated among PI-negative cells unless otherwise indicated. For analysis of expansion localization, cells were manually classified according to the spatial distribution of phase-bright regions as polar, if regions overlapped with the periphery of a cell pole, or nonpolar, if regions existed anywhere that did not make contact with a cell pole.

Imaging was performed on Nikon Ti2-E or Ti-E inverted microscopes equipped with temperature-controlled stages (Okolab) and 100X Plan Apo oil-immersion objectives (NA = 1.45, Ph3) using NIS-Elements software (Nikon). The Ti2-E was equipped with a SOLA LED light engine (Lumencor) and ORCA-Fusion sCMOS camera (Hamamatsu Photonics), and the Ti-E was equipped with an X-Cite 120 LED light source (Lumen Dynamics) and ORCA-ER CCD camera (Hamamatsu Photonics). Fluorescence filter sets were from Chroma Technology.

### Fluorescent peptidoglycan labeling with HADA

A 100 mM stock of the fluorescent D-amino acid HADA was prepared in DMSO. Log-phase cells (OD_600_ = 0.05) were labeled with 0.5 mM HADA for 1 h at 37 °C with shaking in the dark^33^. To coordinate HADA labeling with the onset of periplasmic expansion, the 1-h labeling period was initiated at different times relative to each treatment: immediately before washing cells into HADA-free 1X M9 salts for starvation; 30 min after addition of ciprofloxacin; 20 min before addition of globomycin; or 4 h after addition of mitomycin C. Following labeling, cells were pelleted at 6,010*g* for 1 min and resuspended in 1X M9 salts for starvation experiments or in fresh LB containing the respective antibiotic for antibiotic-treated samples. HADA was omitted from all media during imaging. Samples were immediately spotted onto agarose pads and imaged by phase-contrast and fluorescence microscopy using a DAPI filter set.

### Cell length quantification and microfluidic imaging

Cell lengths were measured from phase-contrast images using the MicrobeJ plugin in FIJI/ImageJ. For time-lapse measurements of relative changes in cell length, images were acquired at 15-s intervals. Rapid changes in cell length following sudden starvation or osmotic shock were measured using a CellASIC ONIX2 microfluidic system (Sigma-Aldrich). Cells were first maintained under continuous flow of LB to establish baseline growth. For starvation experiments, the medium was then switched to 1X M9 salts. For osmotic-shock experiments, the medium was switched to LB supplemented with 500 mM MOPS.

### Periplasmic and cell area measurements

Cells expressing periplasmic mCherry were spotted onto LB agarose pads containing 20 µg/mL globomycin and imaged by phase-contrast and epifluorescence microscopy at 60-s intervals. Individual cells were analyzed using the MicrobeJ plugin in FIJI/ImageJ. Total cell area was determined from phase-contrast images, and the area of the expanded periplasm was determined from the corresponding periplasmic mCherry signal at each time point. Changes in periplasmic area and total cell area were then tracked over time for individual cells.

### Metabolomic extraction and UHPLC-MS analysis

Metabolites were extracted from exponentially growing cultures or from cultures collected 45 min after sudden starvation. Cells were pelleted and quenched in ice-cold methanol:acetonitrile:water (2:2:1, v/v/v), followed by bath sonication for cell lysis. Extracts were stored at –20 °C overnight, thawed, and centrifuged at 14,000*g* for 10 min at 4 °C. Supernatants containing extracted metabolites were collected, and both the supernatants and remaining cell pellets were dried using a vacuum concentrator (SpeedVac, Thermo Fisher Scientific). Dried cell pellets were weighted to determine dry cell mass, which was used to normalize metabolite extracts. Dried metabolite extracts were reconstituted in acetonitrile:water (2:1, v/v) at a volume of 40 µL per mg of dry cell mass. Samples were bath-sonicated at 25 °C for 5 min and vortexed for 1 min; this cycle was repeated twice. Samples were stored at 4 °C overnight and centrifuged at 14,000*g* for 10 min at 4 °C. Supernatants were transferred to LC-MS vials for analysis.

Ultra-high-performance liquid chromatography-mass spectrometry (UHPLC-MS) was performed using a Vanquish Flex UHPLC system coupled to an Orbitrap ID-X Tribrid mass spectrometer (Thermo Fisher Scientific). Polar metabolites were separated on an iHILIC-P Classic column (100 × 2.1 mm, 5 µm; HILICON) equipped with a matching guard column (20 × 2.1 mm, 5 µm). Mobile phase A consisted of water:acetonitrile (95:5. v/v) containing 20 mM ammonium bicarbonate, 2.5 µM medronic acid, and 0.1% ammonium hydroxide; mobile phase B consisted of acetonitrile:water (95:5, v/v). The column was maintained at 45 °C. The gradient was run at 250 µL/min as follows: 90% B from 0 to 1 min; a linear decrease to 35% B from 1 to 12 min; a linear decrease to 25% B from 12 to 12.5 min; 25% B from 12.5 to 14.5 min; and a return to 90% B by 15 min. The column was re-equilibrated at 90% B for 4 min at 400 µL/min followed by 2 min at 250 µL/min. Metabolomic data were processed and annotated using X13CMS^34^, CompoundDiscoverer, and Skyline^35^.

## Acknowledgements

We thank Corey Westfall for assistance with strain construction and Alexander Liu for computational support. This work was supported by NIH grants R35 GM127331345 (to P.A.L.) and DP1 DK147449 (to K.C.H.), NSF grant EF-2125383 (to K.C.H.), an Ignite Interdisciplinary Grant (to A.M.B, P.A.L, K.C., and G.J.P.), and a Howard Schneiderman Fellowship (to A.M.B.).

## Contributions

A.M.B, K.C.H., and P.A.L conceived the project and designed the study. A.M.B., T.L.S., and K.C. performed experiments and analyzed data. G.J.P. and K.C. designed and performed the metabolomics experiments and analyzed the resulting data. A.M.B., K.C.H., and P.A.L. wrote the manuscript with input from all authors.

## Supplementary Figures

**Figure S1:**
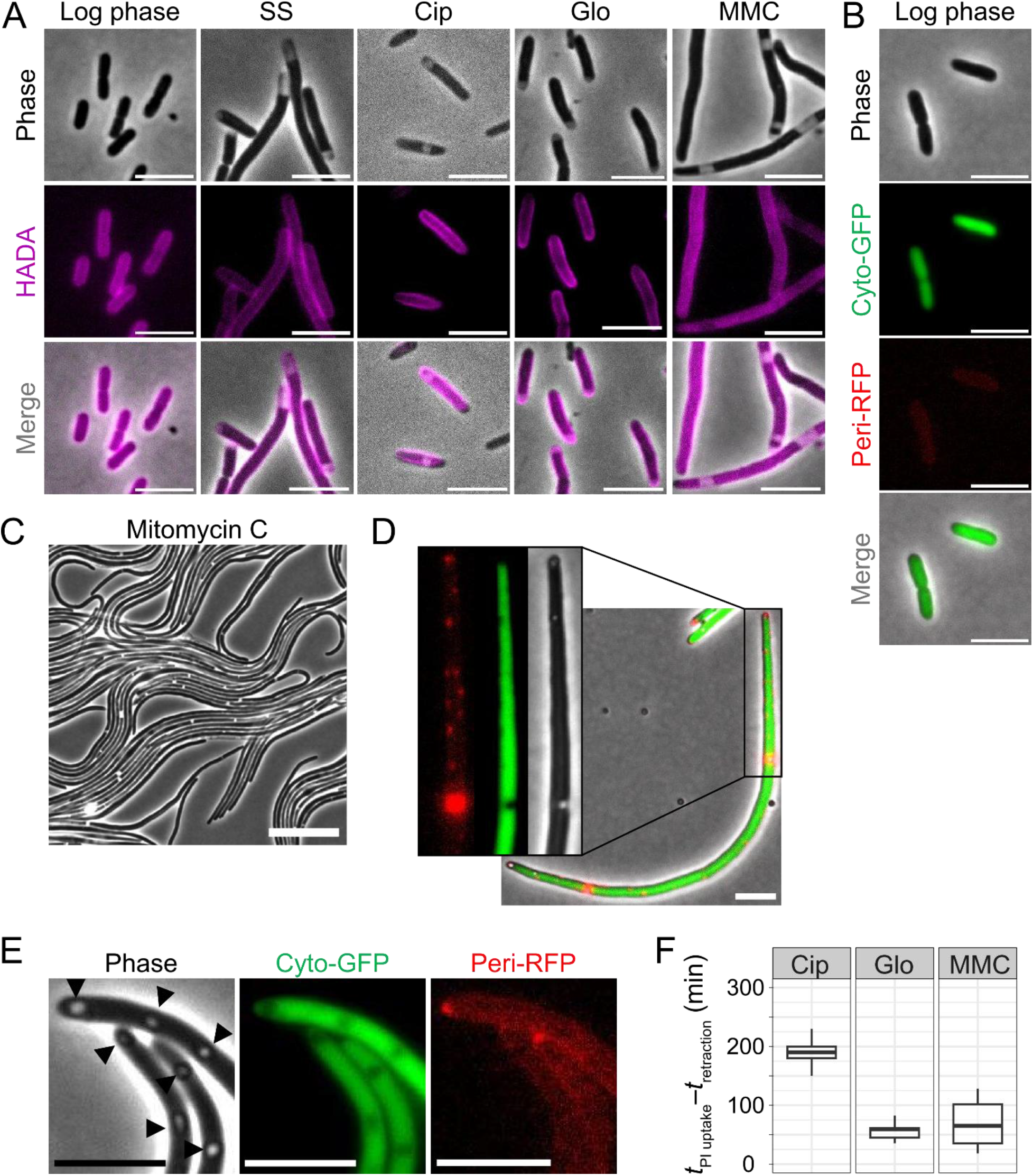
Periplasmic expansion exhibits stress-dependent morphology and dynamics. **A**, Representative phase-contrast and fluorescence images of HADA-labeled cells during exponential growth (log phase), following sudden starvation (SS), or after treatment with ciprofloxacin (Cip), globomycin (Glo), or mitomycin C (MMC). Scale bars, 5 µm. **B**, Representative phase-contrast and fluorescence images of exponentially growing cells expressing cytoplasmic GFP (Cyto-GFP) and periplasmic mCherry (Peri-RFP). Scale bars, 5 µm. **C**, Representative phase-contrast image showing filamentation following mitomycin C treatment. Scale bar, 20 µm. **D**, Representative mitomycin C-treated cell expressing cytoplasmic GFP and periplasmic mCherry. The enlarged region shows multiple small mCherry-enriched regions corresponding to local periplasmic expansions that are not readily apparent by phase-contrast microscopy. Scale bar, 5 µm. **E**, Representative mitomycin C-treated cells containing phase-bright regions (arrowheads) that lack corresponding periplasmic mCherry signal and were therefore excluded from scoring as periplasmic expansion. Scale bars, 5 µm. **F**, Time between the onset of periplasmic expansion and subsequent propidium iodide (PI) uptake in individual ciprofloxacin-, globomycin-, and mitomycin C-treated cells. Boxes indicate the interquartile range (IQR) and median; whiskers extend to 1.5× IQR.

**Figure S2:**
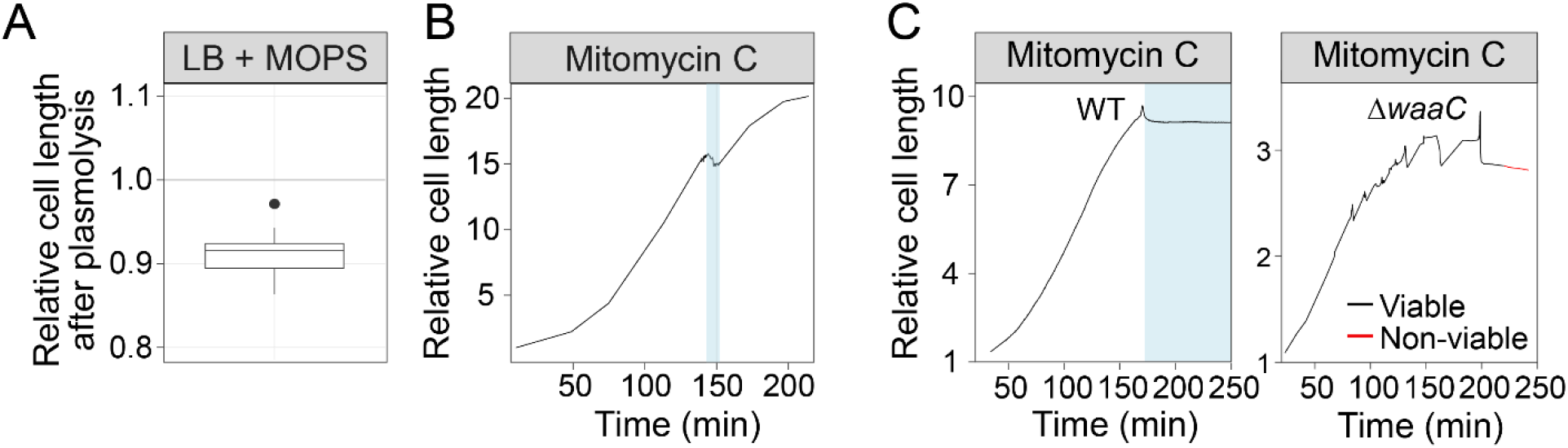
Cell-length changes associated with osmotic shock and mitomycin C treatment. **A**, Relative cell length following plasmolysis induced by osmotic shock with LB supplemented with 500 mM MOPS, compared to cell length before plasmolysis. Box indicates the interquartile range (IQR) and median; whiskers extend to 1.5× IQR; points indicate outliers. **B**, Representative single-cell trace of a mitomycin C-treated cell exhibiting a decrease in cell length coincident with the onset of periplasmic expansion, followed by reversal of periplasmic expansion and resumption of growth. Blue shading indicates the duration of periplasmic expansion. **C**, Representative cell-length traces of mitomycin C-treated BW25113 wild-type (WT) and Δ*waaC* cells. Black and red lines indicates PI-negative and PI-positive states, respectively, and blue shading indicates the period during which periplasmic expansion was visible.

**Figure S3:**
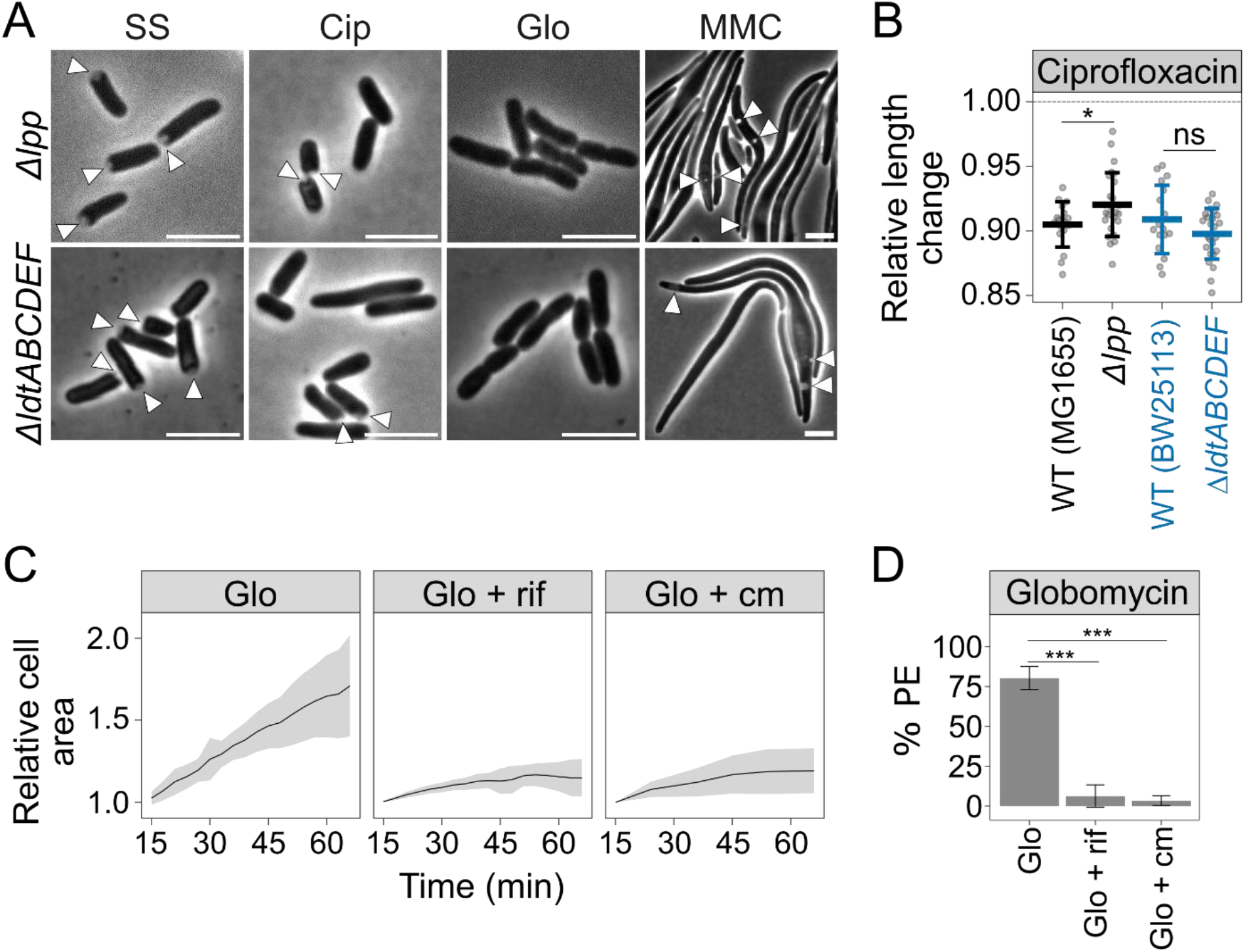
Genetic and physiological requirements for globomycin-induced periplasmic expansion. **A**, Representative phase-contrast images of Δ*lpp* and Δ*ldtABCDEF* cells following sudden starvation (SS) or treatment with ciprofloxacin (Cip), globomycin (Glo), or mitomycin C (MMC). Arrowheads indicate to sites of periplasmic expansion (PE). Scale bars, 5 µm. **B**, Relative cell length following PE compared with cell length before PE in ciprofloxacin-treated Δ*lpp*, and Δ*ldtABCDEF* cells, as compared to their genetic wild-type (WT) parent, MG1655 or BW25113. Points represent individual cells from three biological replicate experiments; bars indicate the overall mean ± SD from all measured cells. Statistical significance was determined using a linear mixed-effects model accounting for biological replicate as a random effect, followed by Dunnett’s multiple-comparison test. ns, not significant (*p* ≥ 0.05); *, *p* < 0.05. **C**, Relative cell area of cells treated with globomycin alone (Glo), globomycin plus rifampicin (Glo + rif), or globomycin plus chloramphenicol (Glo + cm), normalized to cell area at the initial time point. Lines indicate means and shaded regions indicate SD. **D**, Percentage of cells exhibiting PE following treatment with globomycin alone (Glo), globomycin plus rifampicin (Glo + rif), or globomycin plus chloramphenicol (Glo + cm). Data from cells treated with globomycin alone is replotted from Fig. 3A for statistical comparisons. Bars indicate mean ± SD. Statistical significance was determined using a one-way ANOVA followed by Dunnett’s multiple-comparison test. ns, not significant (*p* ≥ 0.05); ***, *p* < 0.001.

**Figure S4:**
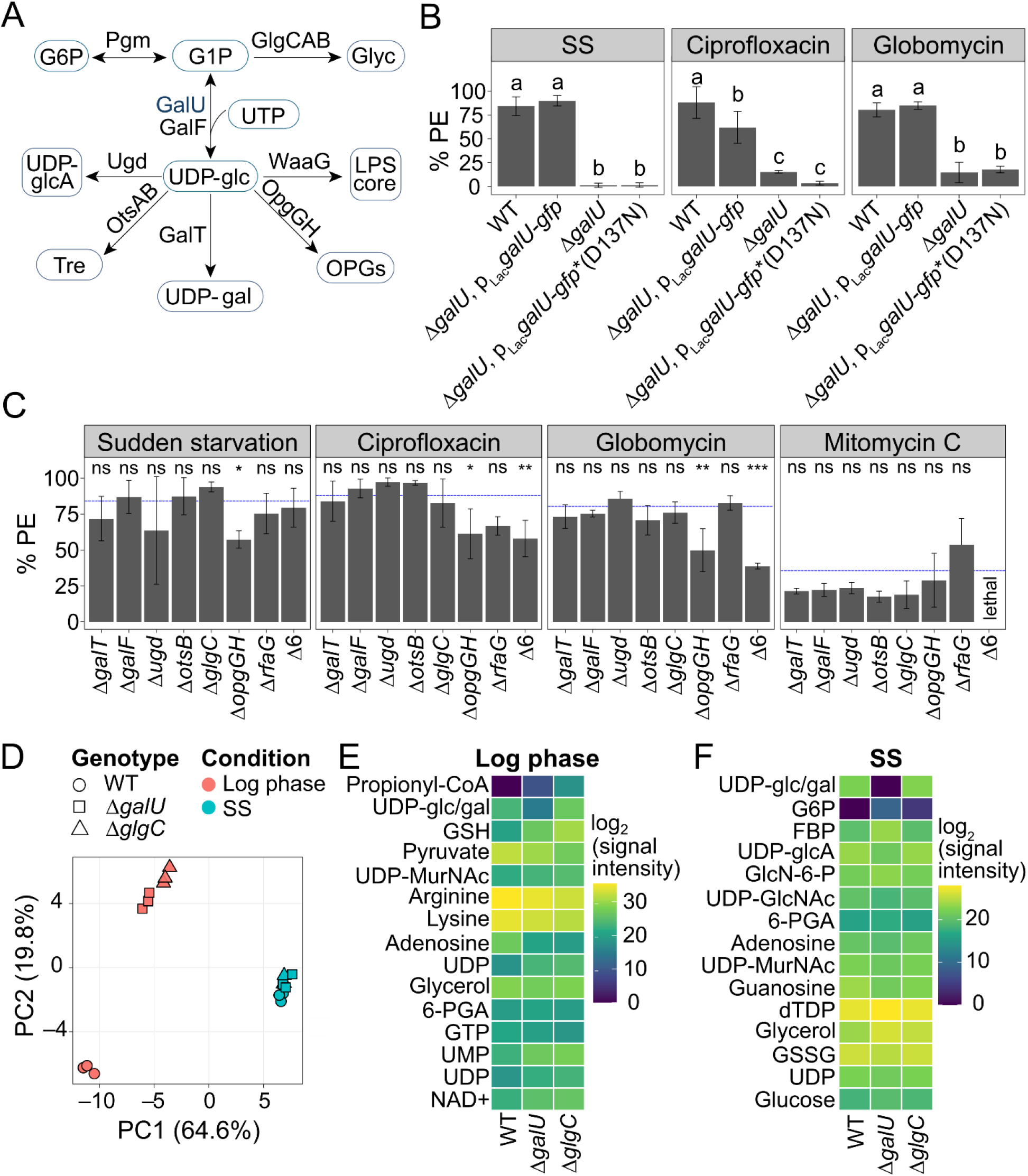
GalU activity is required for robust periplasmic expansion during starvation and treatment with ciprofloxacin or globomycin. **A**, Schematic of UDP-glucose synthesis and selected downstream pathways. G1P, glucose-1-phosphate; G6P, glucose-6-phosphate; UDP-Glc, UDP-glucose; UDP-GlcA, UDP-glucuronic acid; Tre, trehalose; UDP-gal, UDP-galactose; OPGs, osmoregulated periplasmic glucans; Glyc, glycogen. **B**, Wild-type GalU-GFP or catalytically inactive GalU(D137N)-GFP were expressed from a plasmid in the Δ*galU* background in the presence of chloramphenicol for selection. The percentage of cells exhibiting periplasmic expansion (PE) following sudden starvation (SS) or treatment with ciprofloxacin (Cip) or globomycin (Glo) is shown as mean ± SD. WT and Δ*galU* data are replotted from Fig. 5B for statistical comparisons. For each condition, differences among genotypes were evaluated by one-way ANOVA followed by Tukey’s HSD test (*α* = 0.05). Letters (a, b, c) represent statistical significance; genotypes sharing a letter are not significantly different (*p* ≥ 0.05). **C**, Percentage of cells exhibiting PE in mutants disrupting individual or multiple UDP-glucose-dependent pathways following sudden starvation or treatment with ciprofloxacin, globomycin, or mitomycin C. The dotted blue line indicates the corresponding WT mean. Bars indicate mean ± SD. Statistical significance relative to WT within each condition was determined using a one-way ANOVA followed by Dunnett’s multiple-comparison test. ns, not significant (*p* ≥ 0.05); *, *p* < 0.05; **, *p* < 0.01; ***, *p* < 0.001. Δ6, Δ*galT* Δ*ugd* Δ*otsA* Δ*opgGH* Δ*rfaG*. **D**, Principal-component analysis of LC-MS metabolomic profiles from wild-type, Δ*galU*, and Δ*glgC* cells during exponential growth and following sudden starvation. Points represent individual biological replicates; percentages indicate the variance explained by each principal component. **E,F**, Heatmaps showing the 15 metabolites whose abundances most strongly distinguished Δ*galU* from both WT and Δ*glgC* cells during exponential growth (**E**) or following sudden starvation (**F**). Metabolites were ranked by the smaller of the absolute differences in mean log_2_-transformed signal intensity between Δ*galU* and each control genotype. Tile colors indicate mean log_2_-transformed LC-MS signal intensity. UDP-glc/gal, combined UDP-glucose and UDP-galactose signal; GSH, reduced glutathione; UDP-MurNAc, UDP-*N*-acetylmuramic acid; 6-PGA, 6-phosphogluconic acid; UDP-GlcNAc, UDP-*N*-acetylglucosamine; FBP, fructose-bisphosphate; G6P, glucose-6-phosphate; UDP-GlcA, UDP-glucuronic acid; GlcN-6-P, glucosamine-6-phosphate; GSSG, oxidized glutathione.

## Supplementary Tables

**Supplementary Table 1:** Localization of periplasmic expansion under different stress conditions. Cells exhibiting periplasmic expansion were classified according to the location of phase-bright regions as unipolar, bipolar, nonpolar, unipolar + nonpolar, or bipolar + nonpolar. Values indicate the mean percentage of cells in each category ± standard deviation (SD) across *n* independent experiments.

| Condition | Localization | Mean (%) | SD (%) | $n$ |
| --- | --- | --- | --- | --- |
| Sudden starvation | Unipolar | 95.9 | 5.0 | 6 |
|  | Bipolar | 1.8 | 2.4 | 6 |
|  | Nonpolar | 2.0 | 2.6 | 6 |
|  | Unipolar + nonpolar | 0.3 | 0.6 | 6 |
|  | Bipolar + nonpolar | 0.0 | 0.0 | 6 |
| Ciprofloxacin | Unipolar | 76.4 | 8.7 | 3 |
|  | Bipolar | 2.8 | 1.1 | 3 |
|  | Nonpolar | 16.8 | 5.5 | 3 |
|  | Unipolar + nonpolar | 3.6 | 2.7 | 3 |
|  | Bipolar + nonpolar | 0.5 | 0.9 | 3 |
| Globomycin | Unipolar | 90.5 | 3.5 | 5 |
|  | Bipolar | 1.5 | 1.5 | 5 |
|  | Nonpolar | 6.9 | 1.9 | 5 |
|  | Unipolar + nonpolar | 1.1 | 1.5 | 5 |
|  | Bipolar + nonpolar | 0.0 | 0.0 | 5 |
| Mitomycin C | Unipolar | 3.8 | 3.4 | 3 |
|  | Bipolar | 0.0 | 0.0 | 3 |
|  | Nonpolar | 70.9 | 20.9 | 3 |
|  | Unipolar + nonpolar | 21.8 | 15.3 | 3 |
|  | Bipolar + nonpolar | 3.5 | 6.1 | 3 |

**Supplementary Table 2: Raw signal intensity values for metabolomics dataset.** Wild-type (WT), Δ*glgC*, and Δ*galU* cells were harvested during log phase growth and after 45 minutes of sudden starvation. Metabolites were harvested and analyzed with UHPLC-MS. Signal intensity values of *N* = 3 replicates are listed for each metabolite measured.

**Supplementary Table 3:** Strains and plasmids used in this study.

| Strain | Relevant genotype | Plasmid | Source |
| --- | --- | --- | --- |
| AMB11 | MG1655 |  |  |
| KC1193 | MG1655 attHK::Plac-dsbAss-mCherry, pZS21-GFP | pZS21-GFP | <sup>1</sup> |
| PAL2323 | BW25113 |  | <sup>36</sup> |
| PAL5190 | BW25113 *rfaC::kan* |  | <sup>36</sup> |
| AMB34 | MG1655 *galU::frt* |  | This work |
| AMB13 | MG1655 *pgm::kan* |  | This work |
| AMB291 | MG1655 *galT::kan* |  | This work |
| AMB242 | MG1655 *galF::kan* |  | This work |
| AMB186 | MG1655 *ugd::kan* |  | This work |
| AMB144 | MG1655 *otsB::kan* |  | This work |
| AMB61 | MG1655 *glgC::kan* |  | This work |
| AMB33 | MG1655 *opgGH::frt* |  | This work |
| AMB159 | MG1655 *rfaG::kan* |  | This work |
| AMB303 | MG1655 *opgGH::frt, ugd::frt, otsA::frt, rfaG::frt, galT::kan* |  | This work |
| AMB189 | MG1655 *galU::frt, pLac-galU-gfp* | *pLac-galU-gfp* | This work |
| AMB282 | MG1655 *galU::kan, pLac-galU(D137N)-gfp* | *pLac-galU(D137N)-gfp* | This work |
| EAM661 | MG1655 *lpp::kan* |  | This work |
| PAL4407 | BW25113 *ldtA::frt, ldtB::frt, ldtC::frt, ldtD::frt, ldtE::frt, ldtF::frt* |  | <sup>37</sup> |

